# Transparency, a better camouflage than crypsis in cryptically coloured moths

**DOI:** 10.1101/655415

**Authors:** Mónica Arias, Marianne Elias, Christine Andraud, Serge Berthier, Doris Gomez

## Abstract

Predation is a ubiquitous and strong selective pressure on living organisms. Transparency is a predation defence widespread in water but rare on land. Some Lepidoptera display transparent patches combined with already cryptic opaque patches. While transparency has recently been shown to reduce detectability in conspicuous prey, we here test whether transparency decreases detectability in already cryptically-coloured terrestrial prey, by conducting field predation experiments with free avian predators and artificial moths. We monitored and compared survival of a fully opaque grey artificial form (cryptic), a form including transparent windows and a wingless artificial butterfly body. Survival of the transparent forms was similar to that of wingless bodies and higher than that of fully opaque forms, suggesting a reduction of detectability conferred by transparency. This is the first evidence that transparency decreases detectability in cryptic terrestrial prey. Future studies should explore the organisation of transparent and opaque patches on the animal body and their interplay on survival, as well as the costs and other potential benefits associated to transparency on land.

## Introduction

Predation is ubiquitous and exerts a strong selection on living organisms. Often, prey sport cryptic colour patterns that reduce detectability by visual predators, rendering prey hardly distinguishable from their background. Crypsis is achieved if colour patterns are random samples of background colouration [1]. This is challenging, as backgrounds are often complex combinations of elements that can move and that vary in colour and pattern [2]. Background matching is efficient only if all aspects perceived by predators (e.g., colour, brightness, polarization) are matched [2,3]. Given the intimate dependence between their survival and background colouration, cryptic colourations constrain prey movements, and potentially hinder foraging and exploration [2,4]. By contrast, dynamic colour changes [5] or transparency [6] can free prey from background dependency, and improve survival in visually heterogeneous environments. Notably, transparency can minimize detectability against virtually any background [6].

Transparency maximises light transmission, minimising reflection and absorption at all angles and for all wavelengths seen by predators. Transparency is rare on land, with the notable exception of insect wings. Among insects, Lepidoptera (moths and butterflies) typically have opaque wings covered by coloured scales involved in intraspecific communication [7], and antipredator defences such as aposematism [i.e. advertisement of unpalatability, 8], masquerade [i.e. imitation of inedible objects, 9] and camouflage [10]. Yet, wing transparency has evolved independently in multiple Lepidoptera families, often in combination with cryptic colour patterns, as in the Neotropical moth *Neocarnegia basirei* (Saturniidae) or the Malaysian *Carriola ecnomoda* (Erebidae), where transparent wing areas are surrounded by brownish patches. By comparing detection of four real species of conspicuously-coloured butterflies by predators, Arias *et al* [11] recently showed that even if all offered a high visual contrast to predators, fully opaque species were more detectable than species with transparent elements. However, this study did not test whether the observed differences were due to transparency itself or to conspicuous colours covering less surface in transparent species. To rigorously test whether transparency decreases detectability on land, a comparison of the detectability of already cryptic patterns that only differ in the presence/absence of transparent areas is necessary. We here test for the first time whether transparency decreases detectability on already cryptic terrestrial prey, by conducting field predation experiments by free avian predators and using artificial moths.

## Materials and Methods

### Field experiments

We performed predation experiments in May 2018 in southern France, in La Rouvière forest, (43.65°N, 3.64°E) for one 1-week session and at the Montpellier zoo (43.64°N, 3.87°E) for the subsequent two 1-week sessions). Great tits (*Parus major*) and blue tits (*Cyanistes caeruleus*) reported predators in previous similar studies [12,13] are present at both locations. We followed the previously used protocol [12,13] for monitoring artificial prey survival to predation by bird communities. Artificial prey (body and wings) were pinned on green oak *Quercus ilex* tree trunks (>10cm in diameter, with few or no moose cover), every 10m in the forest cover. We put Vaseline and sticky double-faced transparent tape between prey and trunk to avoid ant attacks. We randomly placed artificial moths with edible body, and three types of wings: fully opaque grey wings (C form), wings with grey contour and large transparent windows (T form), and no wings (B form) as a control of body attractiveness (Fig. S1). Prey were disposed vertically and mostly facing north to reduce direct sunlight reflection. We monitored prey survival once per day for the following four consecutive days after placing them on trunks, and removed them afterwards.

### Artificial moths

As in other similar experiments, artificial moths consisted of paper wings and an edible body made of flour and lard [12,14,15]. Triangular shaped moths (triangle 25×36mm, surface of 450mm^2^) did not mimic any real local species, but resembled a generic resting moth (examples in Fig. S1). We designed moths to display poor visual contrast (chromatic and achromatic) against the average trunk colouration of the highly abundant green oaks.

First, we took reflectance spectra of green oak trunk colouration (Fig. S2) and laminated grey paper. We calculated colour and brightness contrasts between paper and trunk as seen by birds. Grey155 was found as rather cryptic (chromatically indistinguishable but lighter than oak trunks, Table S1) but not identical to trunk colouration and was chosen to allow us testing transparency as a crypsis enhancer (see ESM for details). We built the “T” form by cutting two triangular windows (total area of 234 mm^2^) in the laminated grey triangle, and putting a transparent film (3M for inkjet, chosen for its high transparency even in the UV range see ESM, Fig S2) underneath the remaining parts. On top of moth wings, we added an artificial body made from pastry dough (428g flour, 250g lard, and 36g water, following Carrol & Sherratt [16]), dyed in grey by mixing yellow, red and blue food dyes (spectrum in Fig. S2, contrast values in Table S1). Such malleable mixture allowed us to register and distinguish marks made by bird beaks from insect jaws. We finally computed the visual contrasts produced in the eyes of bird predators: C was cryptic (ΔS<1JND, ΔQ≤1.64 JND) and more conspicuous than T and B forms (Table S1).

### Data collection and analysis

During monitoring, we considered artificial moths as attacked by birds when their body showed V-shaped or U-shaped marks, or was missing without signals of invertebrate attacks (i.e. no body scraps left on wings or around the butterfly on the trunk). We removed all remains of artificial moths attacked by birds, but replaced them when attacked by invertebrates or fully missing. Non-attacked prey were considered as censored data. We analysed prey survival using Cox proportional hazard regression [17], with prey form and week and their interaction as factors. By including “week”, the first contrast tests for time and place (by comparing week 1 at La Rouvière, and weeks 2 and 3 at the zoo), while the second contrast test for ‘time’ at the zoo (Table S2). Overall significance was measured using a Wald test. Statistical analyses were performed in R [18] using *survival* package [19].

## Results

In total, we placed 497 artificial moths on trunks, of which 70 were attacked (predation rate: 14.08%). Survival strongly differed between forms (Wald test =24.35, df = 8, p = 0.002): wingless bodies and butterflies with transparent windows were similarly attacked (z = 1.51, p = 0.13) and both were less attacked than opaque butterflies (z = 3.98, p < 0.001, Fig. 1, Table S2). No differences could be detected between attacks registered at La Rouvière and attacks at the zoo (z = -0.04, p = 0.71). At the zoo, more attacks were registered on week 2 (closer to blue and great tit reproduction peak) than on week 3 (z = 0.55, p = 0.003). No interaction between prey form and week was detected (Table S2).

**Figure 1.**
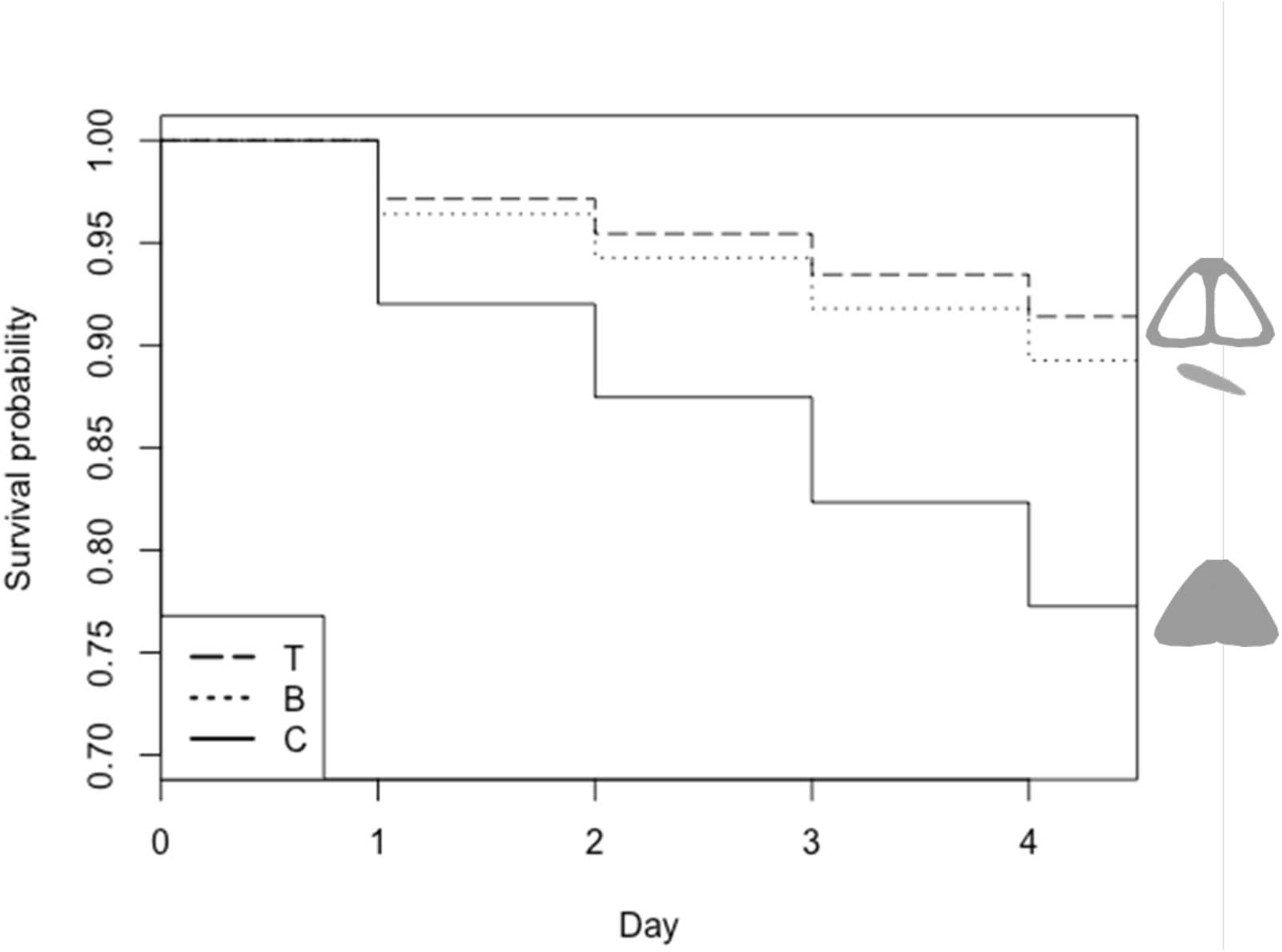
Survival of artificial preys with (T) transparent elements on their wings, (B) bodies without wings, and (C) fully coloured opaque butterflies. Artificial butterflies were placed on tree trunks and monitored for their ‘survival’ every day for 4 days.

## Discussion

Using artificial prey mimicking resting moths with and without transparent elements, we show for the first time that transparency confers survival benefits in already cryptically-coloured terrestrial prey. Transparent butterflies were attacked as little as wingless bodies and less than opaque butterflies, suggesting that transparent windows reduce detection. This study is the first to investigate the benefit value of transparency in cryptic terrestrial prey, and to experimentally isolate the effect of transparency from other aspects (as patch colour or patch size). Whether the position and the size of transparent windows, as well as the intrinsic optical properties of the transparent surface (levels of transmission and reflection briefly explored by [11] and [20]) and its interaction with the ambient light [21] influence transparency efficiency remains untested for terrestrial prey.

Crypsis can incur costs related to thermoregulation [22], intraspecific communication [23], and, more importantly, mobility [2,4], thereby hindering foraging and looking for mates. While costs of transparency in terms of thermoregulation and communication have been unexplored so far, transparency can potentially reduce detectability in virtually all backgrounds, reducing the mobility costs associated to crypsis, and enlarging habitat exploitation as reported for the transparent form of the *Hippolyte obliquimanus* shrimp [24]. However, if camouflage is maximal when including transparency and offers additional benefits in terms of mobility, the low representation of transparency in land, especially in Lepidoptera, is puzzling. As it has been hypothesised for benthic habitats, transparency may be more costly than pigmentation [25]. In Lepidoptera, scales are involved in several physiological adaptations (communication, water repellency, thermoregulation) [7,26,27]. Whether transparent wings may incur communication, hydrophobic or thermal costs remains to be studied to better understand the costs associated to the evolution of transparency on land and explain its rarity.

## Supporting information

ESM

## Acknowledgments

We thank Amandine Aullo, Tiffanie Kortenhoff, Mireia Kohler, Baptiste Laulan, Manon Duty, Samuel Perret and Pablo Giovanni for their major contribution to prey elaboration and fieldwork, and Montpellier zoo for their support. We thank Johanna Mappes and Tom Sherratt for useful comments on the experimental set-up.

## Author contributions

MA, ME, CA, SB and DG designed the study. MA and DG performed the experiments, did the optical measurements, and analysed the data. MA wrote the manuscript with major contributions of ME and DG. All authors approved its final version.

## Accessibility

Data available from Dryad Digital Repository https://datadryad.org/review?doi=doi:10.5061/dryad.82n006f

## Ethical statement

No animal was intentionally harmed during this experiment and most artificial prey were found and collected, avoiding leaving wastes in the forest.

