## Supplementary material for "Transparency, a better camouflage than crypsis in cryptically coloured moths": ESM

**Electronically supplementary material**

### Supplementary Material and methods

#### Artificial prey

We measured reflectance spectra of green oak trunk coloration (120 spectra on various oak barks Fig S1) and of a series of grey colours printed on Sketch Canson paper, laminated with a Polyester Opale Mat 75µm poach. All spectra were measured in spectrometry, using a deuterium halogen lamp (Avalight DHS) emitting over 300-700 nm range including UV to which birds are sensitive [1], a spectrophotometer (Starline Avaspec-2048 L), an optic probe with tip cut at 45° (FC-UV200-2-1.5 x 100, Avantes), and a white diffuse reference (spectralon, WS2).

We then computed the chromatic and achromatic contrasts between paper and trunk as perceived by birds (see below), by using Vorobyev and Osorio's discriminability model [2]. These models reconstruct the difference in colour and brightness between two colour patches (here butterfly and trunk), as seen by a predator under a given light environment. Contrasts are expressed in JNDs (just noticeable differences) and 1 JND is commonly assumed as the threshold below which two colours are indistinguishable.

Bird communities include great and blue tits with UVS vision, and woodpeckers with VS vision. We thus computed visual contrasts for the blue tit as a model for UVS vision [3] and the shearwater as a model for VS vision [4], as its vision is typical of most VS birds. Given that prey items and oak trunks are located under the forest cover, we used a forest shade light environment [5]. We used a Weber fraction of 0.1 for the chromatic response (as reported for Pekin robin *Leiothrix lutea* in [6]) and 0.2 for the brightness response (as the average reported values for known bird species [7]).

We chose the grey155 (R=G=B=155) as close in colour but lighter than trunk coloration for both visual systems (Table S1). This rather cryptic colour was not too cryptic to allow us testing transparency as a crypsis enhancer. On top of butterfly wings, we added an artificial body (428g flour, 250g lard, and 36g water) following Carrol & Sherratt [8], dyed in grey by mixing yellow (10 drops), red (15 drops) and blue (5 drops) food dyes (spectrum in Fig. S2, contrast values in Table S1).

We compared the transmittance of different transparent films to choose the film that transmitted most across the entire range of wavelengths seen by birds (from 300 to 700 nm [4]). We measured transmittance spectra with the same light source and spectrophotometer but with dissociated optic fibres (FC-UV200-2-1.5 x 100, Avantes). Fibers were aligned and the transparent film was held perpendicularly at an equal distance of ~2mm from each fibre. The transparent layer was coated with a transparent mate varnish which increased transparency efficiency by reducing light scattering.

To estimate prey perception, we computed mean chromatic and achromatic contrasts for each prey form by using Vorobyev and Osorio's discriminability model [2] with the aforementioned parameters. We assumed that the color of the transparent area is computed as light going through the wing reflected by the trunk and through the wing again, while for the opaque area, light is reflected on the patch only (as in [9]). Contrasts between background and body, transparent wings or opaque wings, were weighted by the relative surface that either body, opaque and transparent areas covered in each form to compute average contrasts for each form. Surface occupied by the body was measured using GIMP [10]. C was the most achromatically contrasting treatment for both UVS and VS vision systems (see Table S1). All chromatic contrast values were below zero (Table S1).

Supplementary Results

Table S1. Visual contrasts produced by artificial butterflies in predators' eyes

| | UVS vision | | VS vision | |
|---|---|---|---|---|
| Contrast with average oak trunk | Chromatic contrast | Achromatic contrast | Chromatic contrast | Achromatic contrast |
| Grey155 | 0.75 | 2.14 | 0.70 | 2.11 |
| Opaque form (C) | 0.77 | 1.64 | 0.71 | 1.63 |
| Transparent wings (T) | 0.49 | 0.78 | 0.42 | 0.76 |
| Body only (B) | 0.82 | 0.54 | 0.73 | 0.56 |

Values computed with Vorobyev and Osorio's discriminability model for UVS and VS vision are expressed in JNDs (just noticeable differences). The transparent form shows the lowest chromatic contrast and brightness contrast decreases from the opaque form, to the transparent and the body alone forms.

Table S2. Cox regression analyses for attacks on artificial butterflies.

| | Coefficient | Exp. Coef | SE | *z* | *p* |
|---|---|---|---|---|---|
| Form. C > T | 0.69 | 1.99 | 0.17 | 3.98 | <0.001*** |
| Form. B > T | -0.34 | 0.71 | 0.22 | -1.51 | 0.13 |
| Rouvière w1 < Zoo w2-3 | -0.04 | 0.96 | 0.10 | -0.38 | 0.71 |
| Zoo w2 > Zoo w3 | 0.55 | 1.73 | 0.18 | 3.01 | 0.003** |
| Form. C > T : Rouvière w1 < Zoo w2-3 | 0.76 | 1.08 | 0.12 | 0.63 | 0.53 |
| Form. B > T : Rouvière w1 < Zoo w2-3 | -0.05 | 0.95 | 0.15 | -0.31 | 0.76 |
| Form. C > T: Zoo w2 > Zoo w3 | 0.21 | 0.81 | 0.21 | -0.99 | 0.32 |
| Form. B > T: Zoo w2 > Zoo w3 | 0.45 | 1.57 | 0.29 | 1.58 | 0.11 |

Explanatory variables are: form (fully coloured 'C', with transparent windows 'T', and wingless bodies 'B') week contrasts testing for: time and place (La Rouvière week 1 and Zoo weeks 2 and 3 (Rouvière w1 < Zoo w2-3)), and time at the zoo (Zoo w2 > Zoo w3); and their interaction. *z* correspond to the values from the Wald z test used to test for factor significance, Exp. Coeff. to exponentiated coefficients or hazard ratios and SE to standard errors. Symbols: ** $p < 0.01$, *** $p < 0.001$.

a. b. c.

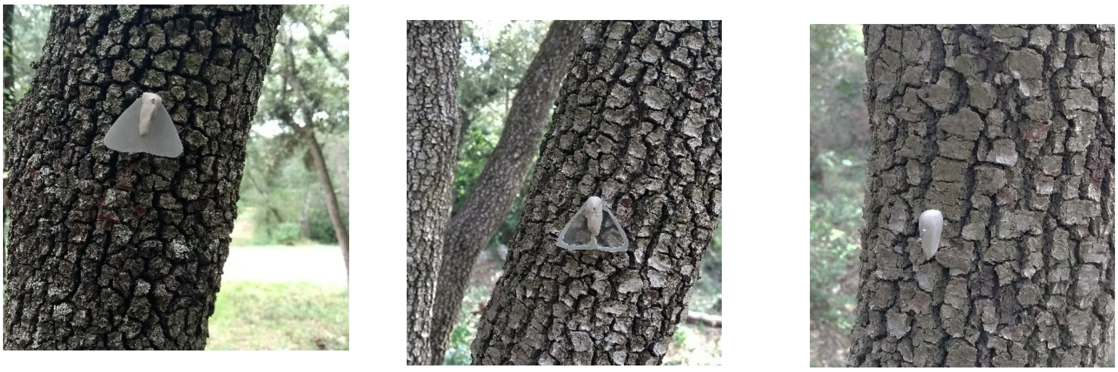

Figure S1. Examples of pictures of artificial prey butterflies placed on oak trunk: a) Coloured ‘C’ form, b) transparent ‘T’ form and c) body ‘B’ form.

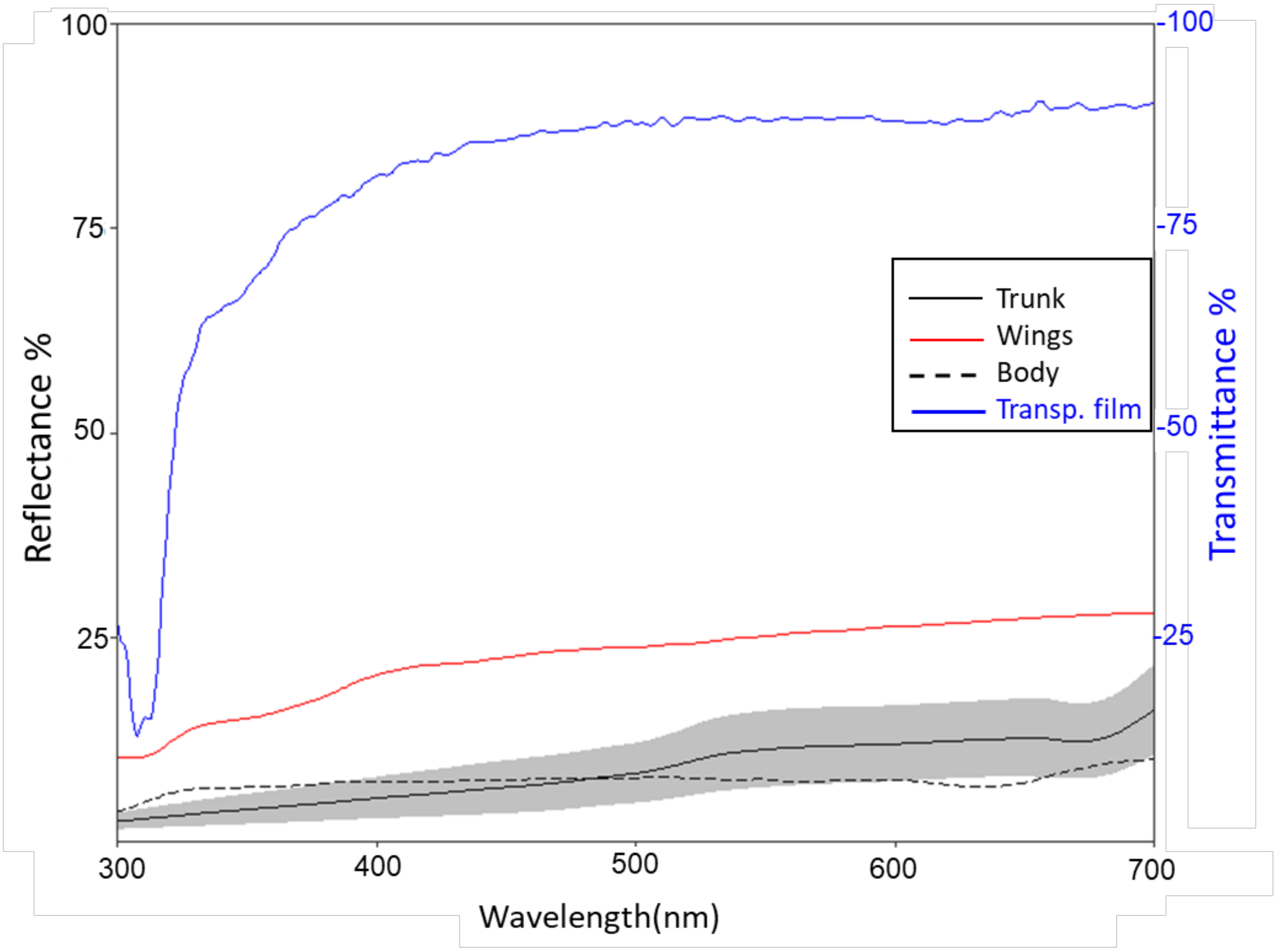


Figure S2. Reflectance of green oak trunks (mean represented by solid line with a confidence interval of ± 1 standard deviation), artificial butterfly bodies (dashed line) and artificial butterfly wings (dotted lines. In blue is the transmittance of transparent films used in the experiment to make transparent windows on prey wings.
